# Single Nanoparticle Tracking Reveals Efficient Long-Distance Undercurrent Transport above Swarming Bacteria

**DOI:** 10.1101/657353

**Authors:** Jingjing Feng, Zexin Zhang, Xiaodong Wen, Jianfeng Xue, Yan He

**Affiliations:** Department of Chemistry, Key Laboratory of Bioorganic Phosphorus Chemistry & Chemical Biology (Ministry of Education), Tsinghua University, Beijing, 100084, China; State and Local Joint Engineering Laboratory for Novel Functional Polymeric Materials, College of Chemistry, Chemical Engineering and Materials Science, Soochow University, Suzhou, 215123, China; Centre for Soft Condensed Matter Physics and Interdisciplinary Research, Soochow University, Suzhou, 215006, China

## Abstract

Flagellated bacteria move collectively in a swirling pattern on agar surfaces immersed in a thin layer of viscous “swarm fluid”, but the role of this fluid in mediating the cooperation of the bacterial population is not well understood. Herein, we use gold nanorods (AuNRs) as single particle tracers to explore the spatiotemporal structure of the swarm fluid. We observed that individual AuNRs are transported in a plane of ~2 μm above the motile cells. They can travel for long distances (>700 μm) in a 2D plane at high speed (often >50 μm^2^/s) without interferences from bacterial movements. The particles are apparently lifted up and transported by collective mixing of the small vortices around bacteria during localized clustering and de-clustering of the motile cells, exhibiting superdiffusive and non-Gaussian characteristics with alternating large-step jumps and confined lingering. Their motions are consistent with the Lévy walk (LW) model, revealing efficient transport flows above swarms. These flows provide obstacle-free highways for long-range material transportations, shed light on how swarming bacteria perform population-level communications, and reveal the essential role of the fluid phase on the emergence of large-scale synergy. This approach is promising for probing complex fluid dynamics and transports in other collective systems.

## Introduction

Bacterial swarming is collective migrations of flagellated cells across agar surfaces with swirling patterns^1–4^. During swarming, fast moving bacteria are trapped in a thin layer of viscous fluid called “swarm fluid”^5–8^. The layer of fluid is only micrometers-thick and even thinner at the swarm edge, and has Reynold number as low as 10^−5^. The swarm fluid can trap surfactants, modify the surface tension of liquids, supports the flagella operation, and carry nutrients or other signaling molecules^9–11^. A fundamental challenge is to understand the relationships between bacteria cells and the fluid medium^5,12,13^. Biologically, bacteria can only “sense” changes in the adjacent fluid environment to coordinate their behavior. For instance, in quorum sensing, the cells respond to the accumulations of signaling molecules dispersed in the fluid through gene regulations^14,15^. The bacteria also react to chemical gradients to control their short-time run lengths through rotating the flagellar^16,17^. Theoretical work such as the Vicsek Model^18,19^ is based on the assumption of collisions and alignments of single bacterium with its neighbors in short-range. To consider hydrodynamic interactions between motile cells in the context of large-scale collective dynamics^5,20–24^, some physical models treated the swarm fluid as a continuum entangling the bacteria phase and fluid together^20,24^, in which the bacterial community are treated as discrete individual self-propelled particles surrounded by an incompressible and inseparable fluid^20,25^.

Yet experimental evidences based on single particle tracking (SPT) have shown that the fluid environment is complicated and heterogeneous. SPT has long been used as a powerful tool to investigate complex system^26,27^. Previous studies have used micro-particles to track the dynamics of bacterial fluid^28,29^. Wu *et al*. revealed an intensive matter transfer flow (rate v= 8 μm/s) in the leading edge of the swarm using 1~2 μm micro-bubbles^30^. Zhang *et al*. found 0.2 μm MgO particles dropped on the liquid-air interface (±20 μm from the swarm front) only diffused normally within a small region (~ 4 μm^2^, v= 0.9 μm/s) as the swarm front approaches^31^. Be’er *et al*. found 0.5 μm MgO particles exhibited superdffusive (v= ~ 9 μm/s) on the upper surface (within 100 μm from the leading edge)^32^. However, since the thickness of the fluid layer is comparable to the dimension of the particles, micron-sized tracers could either have obvious collisions with the motile cell bodies or be trapped at the liquid-air interface, making the tracers’ motions incapable of reflecting the real motions of the pure fluid medium.

Here, using *Bacillus subtilis* as a model swarming system^33^, we introduced 40 × 84 nm gold nanorods (AuNRs) as tracers into the upper region of swarm fluid at near the edge of the swarming bacteria monolayer. Have good photostability and low cytotoxicity^34–36^, plasmonic AuNRs have been used as tracers for long-duration observation in biological studies with high temporal and spatial resolution. We observed that the nanotracers move continuously in 2D at ~2 μm above the bacteria layer, advected by the collective vortices resulting from dynamic clustering of bacteria. They could travel rapidly for a total distance of ~800 μm over the top of hundreds of bacterial cells, while having no direct physical contact with any of them. Their trajectories could be best described by a non-Gaussian superdiffusive model, Lévy walk. Compared to random Brownian motion, Lévy walks could lead to highly efficient long-range transport in complex fluid^37–39^. Therefore, mediated by the swarm fluid, the highly active bacteria community create a dynamically well-organized flow transport network above their own bodies, possibly providing a long-range communication avenue for the population. These discoveries may help to understand the information exchange in the bacterial population and the swarm dynamics. This single nanoparticle tracking method could be potentially applied to probe the fluid dynamics and matter transport in other collective systems.

## Experimental results

### Gold nanorods are lifted above the swarming bacteria

To study bacteria swarming, we chose the wild-type *Bacillus subtilis* 3610 strain as the model system^33^. After ~ 2 h inoculation the bacterial colony begins swarming on the wet agar surface and expands outward in swirling patterns, forming a dendrite-like edge pattern (Figure 1a). The characteristic scale of the swirls is about 10 μm and they last for about 0.2 s. To monitor the bacteria fluid by SPT, we used monodispersed 40 nm × 84 nm AuNRs (Supplementary Fig. 1). They were modified with SH-PEG moieties to make them electrically neutral, so that to prevent their adhesion to the negative charged bacterial cell surface. To introduce the AuNRs into the fluid without causing disturbances, we used a micro-sprayer to atomize the AuNR solution at about 20 mm above the biofilm. By producing an aerosol, the nanotracers naturally fall onto the surfactant fluid surface with minimum interference to the bacterial community. The gravity makes the AuNRs penetrate into the liquid rather than float on the air-liquid interface.

**Figure 1.**
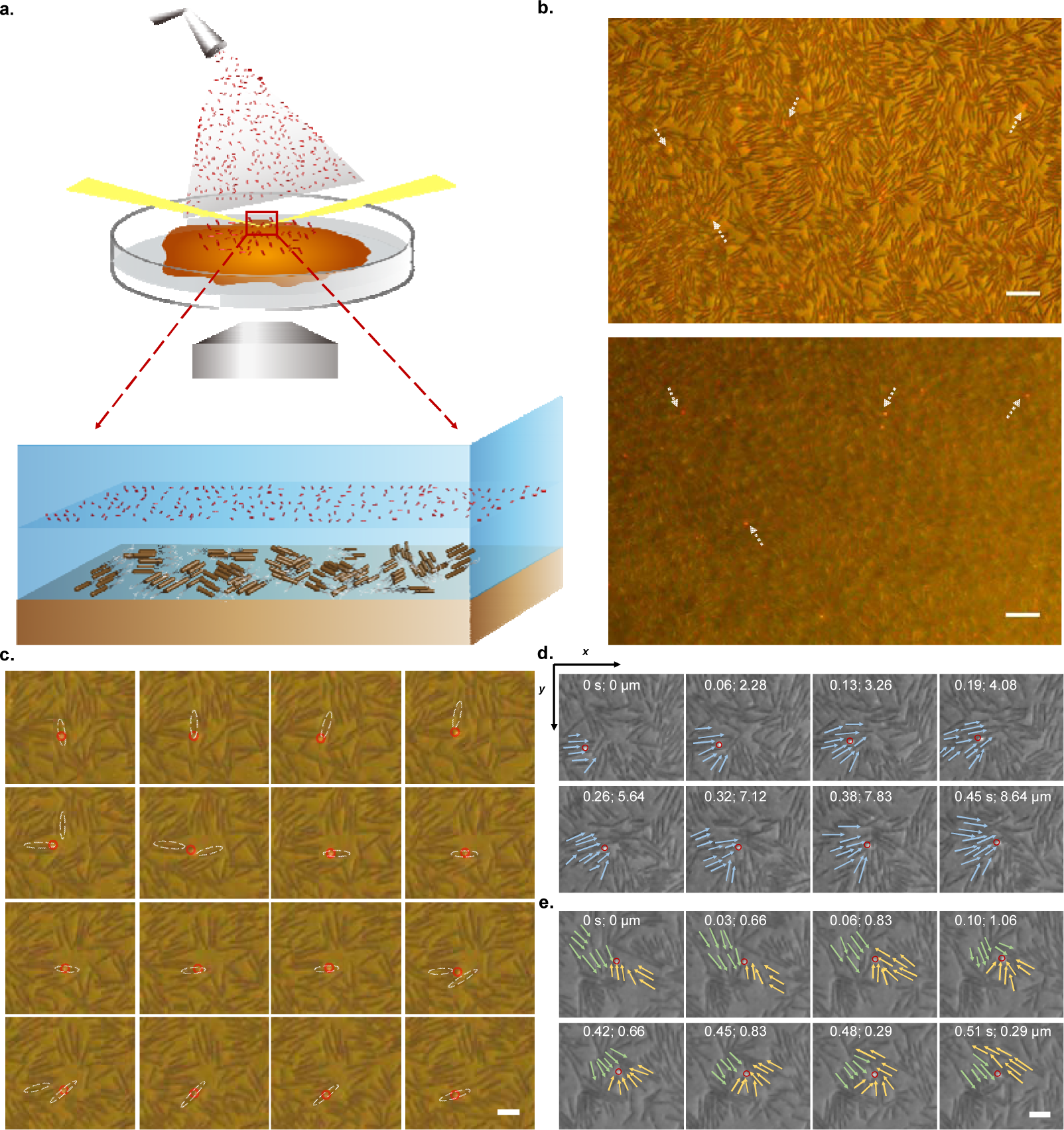
Gold nanorods (AuNRs) float above the swarming bacteria layer. (a) Schematic diagram of the experimental set up as well as the spatial relationship between the observed AuNR tracers and the swarming bacteria. (b) Typical images with the bacteria (upper) and the AuNRs (lower) in focus respectively. The single AuNR are pointed out with a white arrow. The scale bar is 15 μm. (c) A single AuNR crossed over different individual bacterium’s body. The white dash line circles the position of the related bacteria and the red circles mark the position of the single moving nanorod. (d) An AuNR was transferred in one direction by a “vehicle” consisting of a group of aligned bacteria. (e) The motion of an AuNR slowed down when bacteria clusters encounter a “traffic jams” caused by collision. The arrows show the moving direction of the bacteria. The red line circles the nanorod position. The coordinates in (x, y) shows the time (s) and the displacement (μm) of the AuNR. The scale bar is 5 μm.

For imaging, we used an inverted darkfield microscope with a 20X long working distance objective and a color CMOS camera (Supplementary Fig. 2). The probing area is near the edge of the monolayer of swarming bacteria where the cells are most active. Under circular oblique illumination, the broadband absorption of the individual bacteria and the plasmonic scattering from single AuNRs allow them to be imaged simultaneously (Figure 1b). The former appears dark and the latter appears red since the plasmonic scattering maximum of the AuNR is 650 nm. Interestingly, different from the 3D Brownian motion in a passive homogenous solution, i.e. being out-of-focus stochastically, the single AuNRs are seemingly sliding on a 2D plane, with high lateral moving rates but little axial movement (Supplementary Movie 1). We could generally observe dozens of AuNRs in one frame and most of the single AuNRs could be tracked in focus continuously for tens of seconds until they run out of the boundaries of the view field. With the particles keep moving in-and-out, a considerable number of particles can be seen even after 1 hour of their addition.

Close examination of the single AuNR motions indicates that they move just like single bacterium but are relatively independent. They walk, jump or run stochastically among the bacteria dynamic clusters at fast but varying speeds. At any given time, however, the motion of the single AuNRs could hardly be attributed to the physical contact such as dragging, pushing or collision from a single bacterium. Although most bacteria appear to be locally clustered together, many AuNR trajectories are straight and elongated (Supplementary Movie 2) covering several hundred bacteria. Some particles are even able to cross half of the field of view, traveling a net distance of ~150 μm and a total distance of ~ 760 μm. Comparing the sequence of consecutive positions of single AuNRs with nearby single bacteria indicates that the trajectories of nanotracers and cells often cross but do not affect each other (Figure 1c). Therefore, the AuNRs must be transported in a thin 2D layer above the bacteria monolayer. By optical slicing using a reflective laser confocal microscope, the axial distance between the two layers is determined to be ~ 2 μm.

Control experiments indicate that the elevation of the single AuNRs results from the dynamic interactions between the nanotracers and the active bacteria. On one hand, not many tracers could exhibit similar behaviors as the 40 × 84 nm AuNRs. Particles of lower density such as 0.5 μm diameter polystyrene microbeads (Supplementary Fig.1, Movie 3) stayed afloat on the air/liquid interface and were almost motionless. Heavier particles such as 120 nm Au nanospheres (Supplementary Fig.1, Movie 4) swiftly penetrated the entire liquid and bacteria layer, sticking to and even sinking into the agar gel. On the other hand, only the swarming bacteria could raise and drive the nanotracers in the fluid. The AuNRs perform Brownian motions both in the culture medium having moving cells but without swarming (diffusion coefficient D= 3.6 μm^2^/s) and in the culture suspensions where the bacteria are filtered out of the medium (D= 2.5 μm^2^/s) (Supplementary Fig. 3). Moreover, if the bacteria are killed under UV light illumination and stop swarming (Supplementary Movie 5), the AuNRs could only undergo slow Brownian diffusion (D= 0.08 μm^2^/s) in the upper viscous solution, and no fast, long-distance transport could be observed (Supplementary Fig. 3).

### The AuNRs are advected by the collective flow of the swarms

With no physical contact between the bacteria bodies and the nanotracers, the driving force for the AuNR moving in a 2D fluid layer above the colony can only come from the advection that is generated by the collective fluidic flow from the swirling cell populations. The bacterial swarm fluid has low Reynolds number, hence the motion of the particles almost completely depends on the local force^7^. Since the AuNRs appear moving stochastically, the local flow fields must be spatially heterogeneous and change frequently, which can only result from the dynamic clustering of multiple bacteria near the particle. Zhang et. al. has reported that bacterial colonies could spontaneously form tightly packed clusters^40^. Within the spatial scale of the clusters, bacterial movements are strongly coordinated, manifesting as a high degree of correlation between cell body orientation, direction of movement, and velocity. Moreover, the dynamic clusters could evolve, collide, fuse and then re-split into small clusters. Since the axial rotation of bacterium and its flagella swinging would generate a small vortex around its body, the dynamic clustering of multiple bacteria would inevitably result in collision, fusion and splitting of their small vortices, and provide the forces to drive the AuNRs. Therefore, rather than being carried by individual bacteria, the AuNRs are “hitchhiking” on top of dynamically formed bacteria clusters (Figure 1d, e). Notably, these cluster “vehicles” are not stable or long lasting. Whenever encountering “traffic jam”, a cluster will swiftly split and recombine with other individual cells. Meanwhile, the AuNR “passenger” will stop transiently and being transferred to another new “vehicle”, leading to their speed or direction variation. Hence, the transport of the single AuNRs is the result of synergistic effect of local bacterial vortices and is realized through heterogeneous hydrodynamic flow above the bacterial layer.

Particle image velocity (PIV) analyses^41^ were performed to qualitatively investigate the relationships between the two-dimensional AuNR motions and the swirls created by the swarms. From the local orientation and distribution patterns of the flow field velocity vectors, we noticed two localized, instant flow field patterns associated with the nanotracers’ motions. One is the “sticking mode” (Mode A, Figure 2a), where the nanotracers are trapped temporarily or “lingering” in a small area. The magnitude of the velocity vectors of the flow field at the AuNR position is close to 0, resulting from that the rotation of the bacterial swirls around the particle cancel out each other. The other is the “jetting mode” (Mode B, Figure 2b), where the nanotracers “jump” large distances in regions between the sticking area. The sum of the flow field velocity vectors and the joint forces of the adjacent bacterial swirls all point to the same direction. Sequential evolution of the velocity vector patterns indicates that the motion of the single AuNRs is alternating rapidly between the two states (Supplementary Movie 6, 7). The “jump-and-linger” mode of the nanotracers appears similar to the “run-and-tumble” mode of the bacterium, making the single AuNRs behave just like active single bacteria^42^, though with different underlying mechanisms.

**Figure 2.**
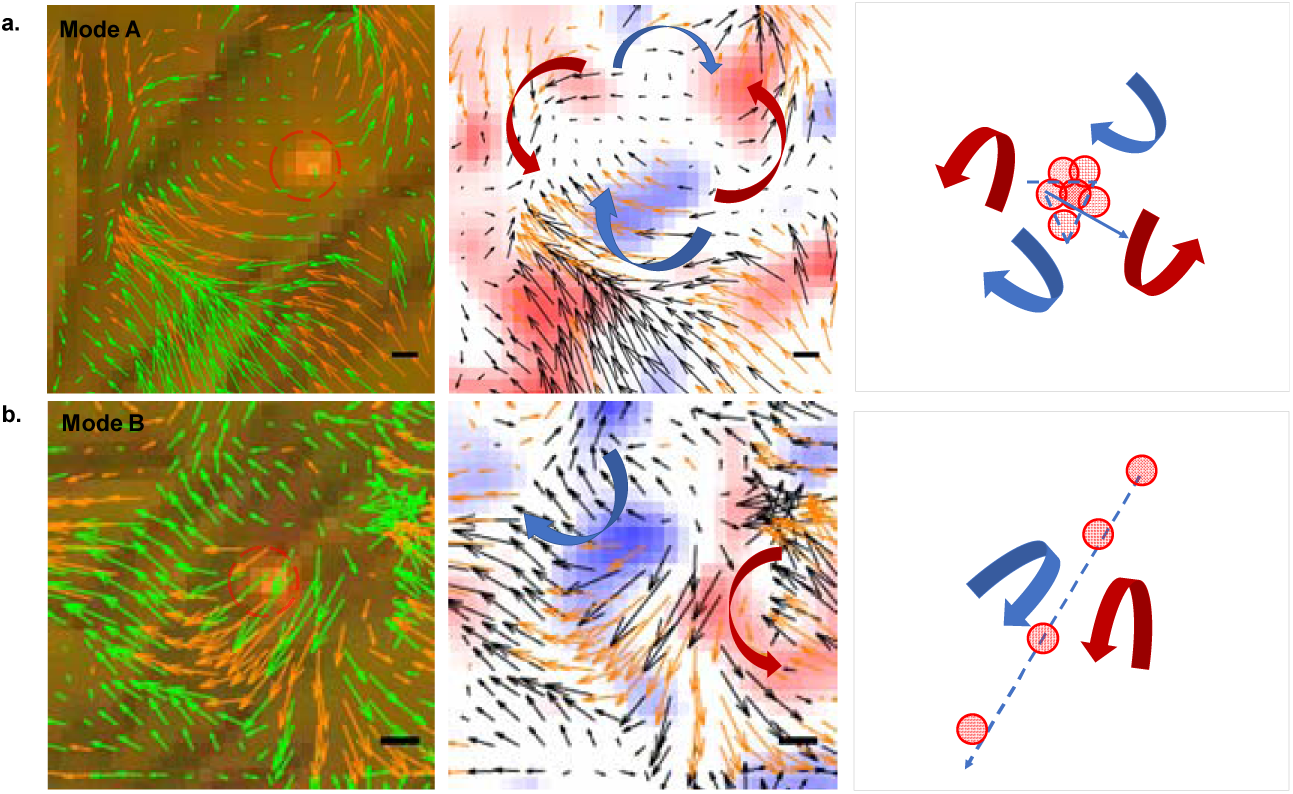
Nanotracer motion states are related to the local flow field patterns. Schematic diagram of (a) the sticking mode and (b) the jetting mode of local flow field patterns. Left panels: the particle image velocity (PIV) vectors overlaid on the raw image. The long dark brown rod is the bacteria and the red-circled spot is the single AuNR. The velocity vectors of PIV are shown in green arrows. The orange arrows are the interpolations. Middle panels: the vorticity color mapping overlaid on the PIV vectors. The velocity vectors of PIV are shown in black and orange arrows. The blue shades refer to the clockwise spin and the red shades refer to the counter clockwise spin. The scale bar is 2.5 μm. Right panels: the red point indicates the position of the particle and the blue line is its trajectory path. In this whole figure, the blue arrow stands for the clockwise flow direction and the red arrow stands for the counter clockwise flow.

### The motions of AuNRs are superdiffusive and non-Gaussian

The phenomenological observation of the AuNR transports in the fluid layer above the bacteria community is further verified through single particle tracking (SPT) analysis. Hundreds of AuNR trajectories are obtained with time duration ranging from 0.02 s to 40 s and traveling distances ranging from 0.3 μm to 800 μm. To get the basic features of the movement, the speed, step size, the motion direction and the difference in motion direction of the AuNRs are examined. We find that the AuNRs move at a speed about 16 μm/s in average (range from 0.03 μm/s to 146 μm/s) and the average step size is about 1.5 μm (0.016 s). As can be seen from Figure 3a and 3b, the single AuNRs at any given time could randomly point to any direction, but the distribution of the variation of the particle directions over a constant time period Δt is centered at zero degree from Δt = 1 to Δt = 60 (0.016 ~ 0.96 s), indicating that the particles tend to keep their directions within short time durations. The mean square displacement (MSD) analyses show that the AuNRs are performing superdiffusion with the ensemble-averaged scaling exponent α of 1.38 and a diffusion coefficient of ~ 21 μm^2^/s (Figure 3c), nearly 300 times larger than normal diffusion in the pure surfactant liquid. The probability density functions (PDFs) show that the distributions of the scaled displacements at different time intervals deviate from the Gaussian. They are both arrow-headed (with a large number of displacements close to zero) and heavy-tailed (decaying more slowly than Gaussian at long displacements) (Figure 3d), suggesting a high probability for the AuNRs to move either at very small steps or at large steps.

**Figure 3.**
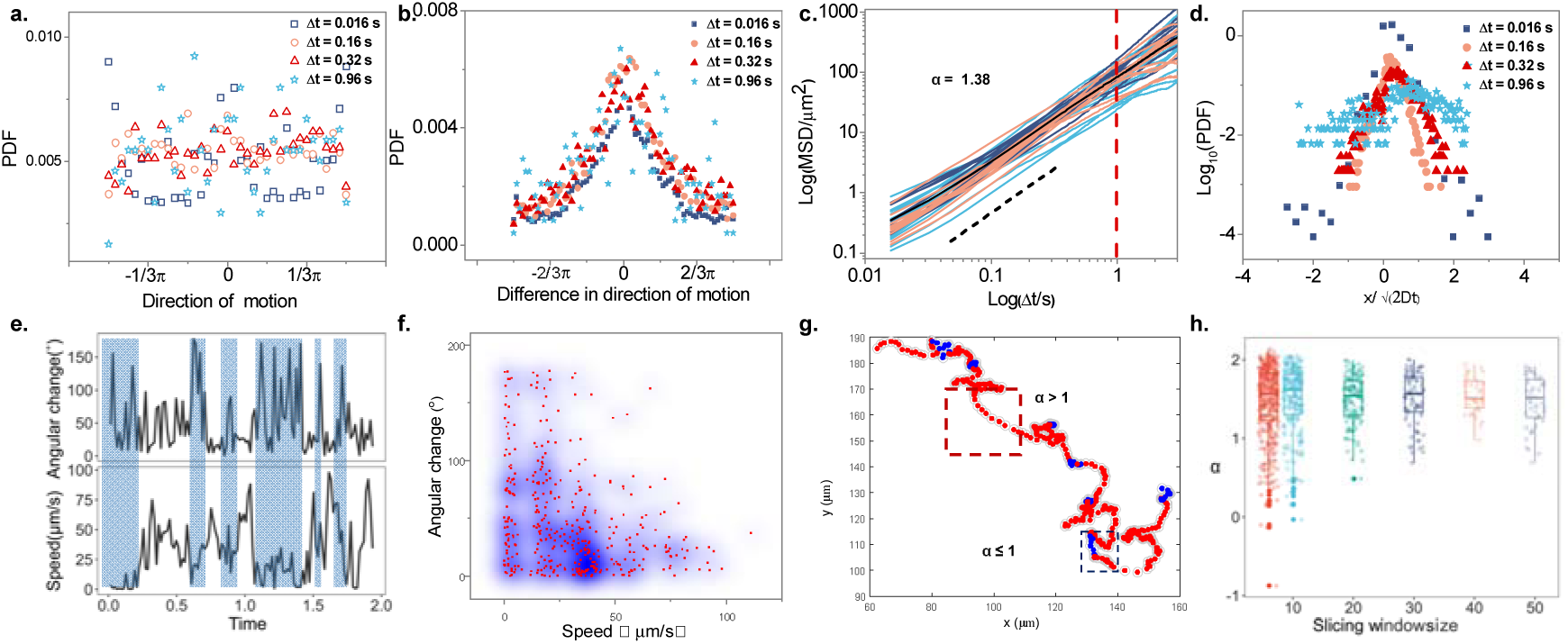
Single particle tracking analysis of the gold nanorods. (a) Probability density function (PDF) of the direction of motion of single AuNRs with different time intervals (Δ t = 0.016, 0.16, 0.32, 0.96 s). (b) PDF of the difference in motion directions of single AuNRs. (c) Mean squared displacements (MSDs) of multiple single AuNR trajectories. Alpha α =1.38 showed by black dash line is the fitted slope of averaged MSD. (d) PDF of the scaled displacements of AuNRs. The y axis is in logarithmic scale. Note that the data in (a, b, d) are calculated by averaging about 57,400 frames. (e) Time-lapsed angular change and the speed of a random sampled trajectory showing the transient motion states between the jump and Brownian-like lingering steps. The rectangles filled with light blue indicate the weak negative correlation between them. (f) The bivariate scattering of the angular change and the speed of the sampled trajectory in (e) with a weak correlation coefficient of −0.33. The blue shades stand for the density. (g) Heat map of the local scaling parameter α showing the presence of both super-diffusive motion (red, α > 1) and the Brownian motion (blue, α ≤ 1) in a single trajectory. (h) Boxplot added with the scatter of scaling parameter α against the slicing window size.

Careful observations of representative AuNR trajectories indicate that there is a weak negative correlation (with a coefficient of −0.33) between the local speed of the particle and the variation degree of the particle direction (Figure 3e, f). The particles tend to keep the original direction when moving fast and change their directions of motion more frequently when the movements slow down, leading to the trajectory appears to be made up of many large jumps with a lot of short pauses in between. To identify the local regions with different dynamic properties, the instant motion states in the trajectory are revealed by trajectory segmentation using temporal slicing windows. By calculating the local scaling exponent from the MSDs, we found that there exist two states, superdiffusive state (jumps) with *α* > 1 and Brownian state (lingering) with *α* ≤ 1 within the same trajectory (Figure 3g). To qualitatively estimate the time scale of the motion states, we tune the size of temporal slicing windows. The sub-trajectories are mostly superdiffusive when the window size is longer than 10 frames. However, when the window size is shortened to 10 frames or less, the local α values exhibits the largest variation (Figure 3h) in the range of *α* ≤ 1, indicating that the Brownian-like movement is more prominent in a time window of less than 0.2 s, which is consistent with the characteristic time of the bacterial swirls.

### Long-range transportations of nanotracers can be described by Lévy walk

Taken together, from both PIV and SPT analysis, we can develop such a picture: during bacteria swarming, individual bacteria create small vortices around their bodies in the surfactant fluid; the coordination of local vortices creates a spatiotemporally heterogeneous flow field above the bacterial layer, allowing both advected rapid transporting and elevated transient trapping of single nanotracers. Interestingly, while the alternating frequency of the AuNR motion state, or the characteristic time of the bacterial cluster formation and dispersion, is not likely to be greater than ~ 0.2 s, the particles have a tendency to keep their walking direction much longer than that. That is, the stochastic switches between the rapid jump and Brownian rest of AuNRs eventually result in their transportation with continually reinforcing directionality.

It has been reported that such directionality persistence, characterized by alternating jump and lingering with superdiffusive and non-Gaussian statistics, often fits with the Lévy process. In particular, the Lévy walk (LW) model describes that the particles walk large distances at a constant speed with local turnings steps. The walking distances mathematically converge to the power-law distribution with no characteristic scales^37^. To test if the single AuNRs were doing LW, “turning points” in trajectories are defined with an instant turning angle larger than a certain threshold^43^ (Supplementary Fig. 4, Note 1). Figure 4a shows a typical long trajectory with a total time of ~ 40 s and total travel distance of 760 μm marked with turning points in an angular speed threshold of 60 rad/s. The distances between two consequent turns are considered as a “flight”. We found that the particle moves at a constant speed of ~16 μm/s through the cumulated length of “flights” as a function of time (Figure 4b). The obtained flight-lengths can be fitted with a power law distribution in the tail by using maximum likelihood estimation. In addition, when choosing different turning-angle-threshold (30, 45, 60 rad/s), the power law index has a variation from −2.38 ~ −2.69 (Figure 4c, Supplementary Note 2). The possibility of matching exponential decays is ruled out by comparing with the Akaike’s information criterion weights^44^. The power-law model is favored with the AIC weights of 0.99. Another evidence is that the power law tail of the velocity correlation function decays as Δ *t*^−δ^ with δ = 0.50 in average^45–47^ (Figure 4d, Supplementary Note 3). According to the previous study, *δ* should equal to 2 − α, when α = 1.38 is the scaling superdiffusive exponent calculated from MSD. Therefore, we can conclude that the motion behavior of the AuNRs is consistent with the LW model. It has been reported that the motion of single bacteria in *B. subtilis* colony is also LW, however, the statistics is slight different^48^. The superdiffusion exponent of the active bacteria is 1.66, indicating more directionality than the nanotracers. Moreover, to get the power law index value of −2.5, the turning-angle-thresholds of the bacteria (5, 10, 15 rad/s) are much smaller than those of the nanotracers, indicating the AuNRs were making turns more dramatically. Therefore, the LW behavior of the AuNRs is similar to the single bacteria motions in the swarm but statistically different.

**Figure 4.**
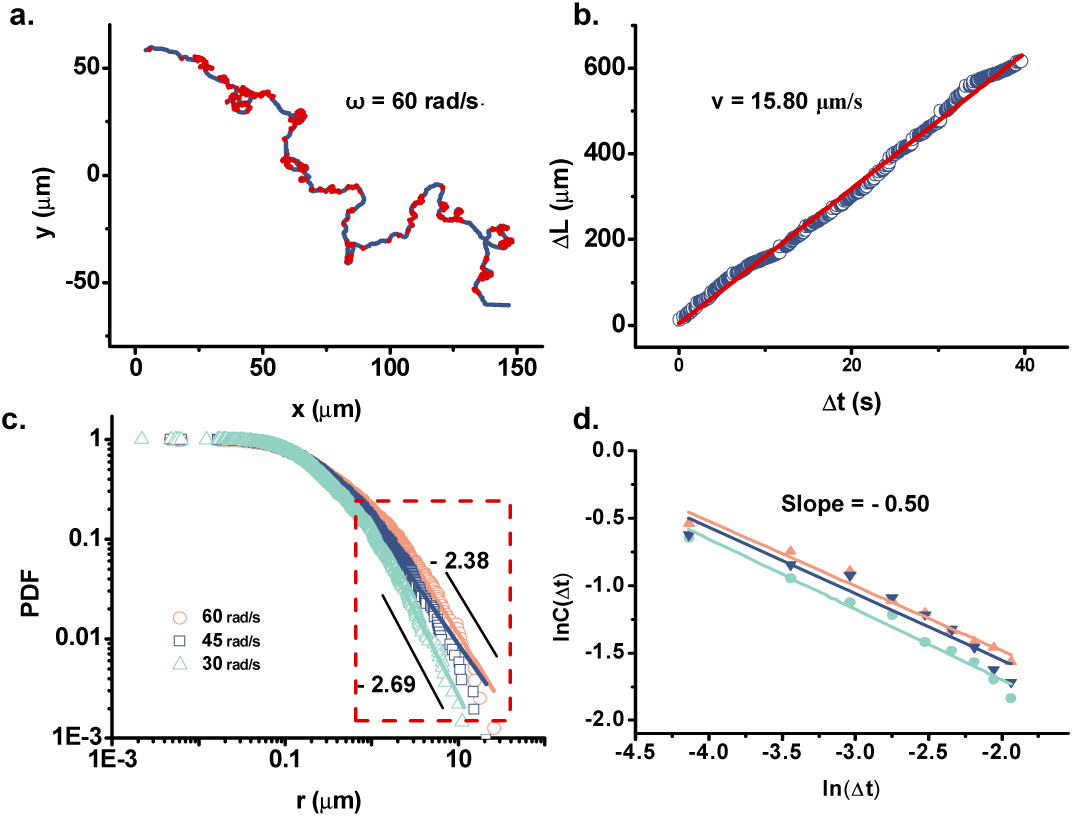
The motions of AuNRs have Lévy walk statistics. (a) The typical trajectory of a single AuNR with turning points marked by red with a threshold angle of 60 rad/s. (b) The cumulated distance ΔL of flights as a function of time. The red line is the linear fitting showing a slope of 15.8 μm / s. (c) Log-log plot of the length distributions of flights against distance showing the power law statistics of turn to turn flight-lengths of the AuNRs. The range of the power law tail is determined by the quality of the fit (see details in the Supplementary Note), highlighted by the red dash rectangle. The fitted slope of the power-law tail is from −2.38 ~ −2.69 with different turning angle thresholds (30, 45 and 60 rad/s). (d) The linear fitting of the natural logarithm plot of the power law tail of the velocity autocorrelation function C(t) as a function of Δt. The color coding stands for different trajectories that collapse well.

### Discussion and Summery

In bacteria swarming systems, how the multicellular collective patterns emerge from a unicellular structure and how information exchange is performed in a swiftly proliferating and expanding population are still a critical issue to explore. With continuous short-range collisions and alignments, it is hard to imagine the existence of certain special “messenger” bacteria capable of carrying and spreading the information rapidly across a wide space. It must be the viscous fluid environment of the swarms that not only enables the self-propelled bacterial cells to spin their flagella to generate thrusts, but also serves as the medium to carry the nutrients and signals from one location to the other in the colony, e.g. the signaling molecules involved in quorum sensing. A previous study deposited MgO microparticles on the surface of the swarm fluid, and superdiffusive trajectories were obtained^32^. Nevertheless, probably due to the large size, no long-range transportation of the microtracers was observed that was associated with the coordination mechanism at large spatial scales.

In this work, the nanotracers are far smaller than single bacteria in size and are lifted above the swarming bacteria. They are neither being pushed by nor “riding” on any single bacterium. Rather, the AuNRs are traveling in 2D in the surfactant fluid layer due to the local synergy of the vortices around the bacteria, and exhibit non-Gaussian superdiffusive characteristics with alternating jumps and lingering, consistent with LW statistics (Figure 5a, b). Under the Lévy motion mode, the AuNR tracers appear no longer passive and could actively and efficiently travel over long distances across multiple bacterial clusters (Figure 5b). Compared with the LW statistics of individual bacteria showed by Ariel *et al*.^48,49^, the LW of single AuNRs have smaller super-diffusion exponents and larger truncating angular speeds, indicating that the AuNRs are moving more random and less directed than the bacteria in the swarms. This could be attributed to the fact that the bacteria are consuming energy and driven by active flagella rotations, but the AuNRs are passively transported by the swarm fluid. Moreover, the bacteria and the AuNRs are spatially divided into two phases with different environments. The single bacteria move in densely packed populations and often encounter physical hindrance from others over short distance. However, the AuNRs are traveling in an obstacle-free fluid layer above the motile cells. The two similar but different LW transports suggest that the collective movement of high-density bacteria cells produced simultaneously two distinct biological mechanisms. The LWs of the single bacteria may have advantages in foraging or avoiding the hazards, which are associated with continuous adaptation to local environments or even “conflict” with neighbors^48,49^. On the other hand, the LWs of AuNR tracers reveal efficient, long-range transportations in the upper fluid medium, which could facilitate communications such as circulations of metabolites and nutrients, fluid mixings for oxygen dissolution, and transports of signaling molecules over long distances. In other words, the two LWs represent the simultaneous near-range competitions and large-scale cooperation between individuals within the same bacteria community, respectively. Interestingly, from the viewpoint of a single bacterium, it should only be able to sense the change of its surroundings and develop its own path through the gaps between neighbors, but would not “foresee” the formation of LW “highways” for efficient transportations and collaborations. Nevertheless, the collective motions of all the individuals lead to the emergence of a long-range communication LW network that “link” their activities altogether.

**Figure 5.**
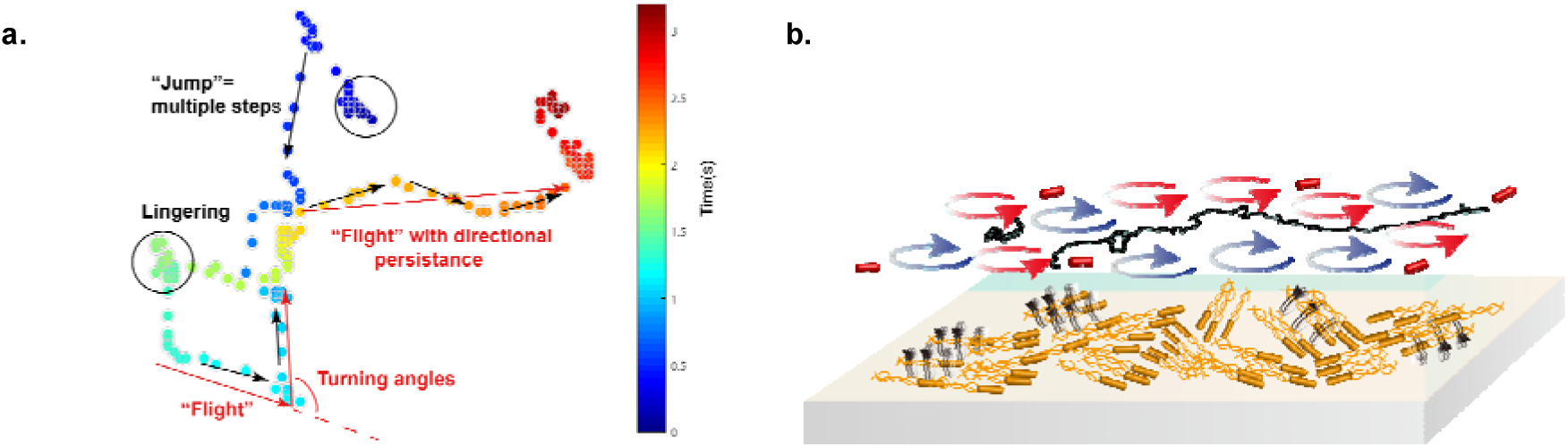
The AuNRs are transported in long range above the swarming bacteria layer by the coordination of the bacteria vortices. (a) Schematic diagram of a representative time-elapsed trajectory of AuNR having jumps, lingering and Lévy walk “flights” with a directional persistence. (b) Schematic diagram illustrates the long-range transportation of the AuNRs by the cooperation of swarming bacteria vortices. The black arrows stand for vortices around the bacteria. The blue and red circles with arrows stand for the vortices in the fluid layer. The black line draws the trajectory of a single AuNR transported by the living bacterial fluids.

Lévy patterns have been identified in a wide range of organisms, from cells, birds, to even human hunters^37,44,50^. Most studies consider LW an optimal search strategy being evolved during adaptive interactions with heterogeneous environments. The LW of passive AuNRs, however, is the result of collective local mixing of vortices around the swarming bacterial bodies. Since the vortices have no clear physical boundaries and are short-lived due to collisions and mixings with each other, our results support a recent hypothesis that the LW can emerge spontaneously and naturally from innocuous responses to the surroundings^51^. Since individual bacteria are also performing LW, to our knowledge, this is the first report of a second Lévy network in a complex living system. The collective behaviors of bacteria produce fluid flows that are physically and functionally separated from the cell bodies. The coincidences and mutual reinforcements of two Lévy networks, possibly one for optimizing the near-range foraging and the other for optimizing the long-range communications, apparently lead to higher-level surviving and adaptation to the environment. Similar mechanisms may constitute the underlying physics of the emergence of large-scale coordination in nature such as flocking of birds, schooling of fishes and collective spreading of cancer cells. Nevertheless, by using nanoparticle tracers, we expect that new collective properties on bacterial and other collective motion systems could be discovered.

## Experimental methods

### Bacteria strain and colony growth

Wild-type *B. subtilis* strain 3610 (purchased from China General Microbiological Culture Collection center) is a gram-positive bacterium with a rod-shaped body and multiple flagella, which generates a propelling force in the direction of its body. They swim with a mean speed about 10 ~ 30 μm/s in a thin liquid film about micrometers thick on the substrate. In our experiments, the bacteria have mean dimensions of 0.7 μm × 4 μm. Colonies were cultured on soft LB agar substrates (10 g/L tryptone, 5 g/L yeast extract, 0.5% agar, all purchased from Bacto Difco, 5 g/L NaCl, purchased from Beijing Chemical Works). For inoculation, 5 μL of *B. subtilis* overnight culture (37 °C, 200 rpm, OD 650 = 0.5) is placed on the center of the agar. The inoculated gel is stored in an incubator at 30 °C. After a lag time of 2 h, the colony starts to expand outward with a speed about 1 cm / hour. After about 2 h, when a colony reached a radius about 2 cm, it could be observed under microscope at the edge of colony where the bacteria almost swarms in a monolayer.

### Tracer materials and the microspray technology

PEG-modified 40×84 nm gold nanorods were purchased from NanoSeedz and were diluted 20 times before use. Carboxyl modified polystyrene microspheres (0.5 μm) were purchased from Dae technique. Gold nanospheres having size about 120 nm were synthesized using a method reported previously^52^. For microspray without disturbing the bacteria colony, we added 50 μL of the chosen particle solution into a high-frequency vibrating atomizer (5W, 5V) and then sprayed it out to a direction parallel to the surface of the colony. The small microdroplets would slowly fall onto the surface of the colony. After that, we immediately took the specimen to the microscope for observation.

### Imaging technique and data analysis

The specimen was placed under an inverted microscope (Nikon Ellipse Ti-U) equipped with a 100 W halogen tungsten lamp, a dark-field air condenser, a 20X long working distance objective lens and an Olympus DP74 color CMOS camera with imaging rate of 60 fps. To localize the precise z-position of the AuNR nanotracer layer relative to the bacteria layer, a confocal microscope equipped with a 20X long working distance objective lens and a 647 nm laser (Nikon A1) was utilized. Single nanoparticle tracking was performed using *Image J* and a code written in *IDL*. Other data analysis was performed using MATLAB and Origin. The segmentation of trajectory using the “angle method”^43^, the quantification of its power-law characteristics, and the evaluation of the power-law model are described in detail in the Supplementary notes. The flow field was measured by using a GUI-based software called PIVlab based on discrete Fourier transform (DFT)^53^.

## Acknowledgements

The authors are grateful to Dr. Shuchun Liu (Institute of Microbiology, Chinese Academy of Sciences) for providing the bacteria strains. This work was supported by the National Natural Science Foundation of China with grant numbers of 21425519, 91853105 and 11574222.

## Author contributions

Y. H. designed the experiments and wrote the paper; J. F. designed and performed the experiments, analyzed the data, and wrote the paper; Z. Z. analyzed the data and wrote the paper; X. W. performed the experiments and analyzed the data. J. X. analyzed the data.

## Conflict of interest statement

The authors declare no competing interests.

## Supplementary notes

### Supplementary Note 1: The angle method

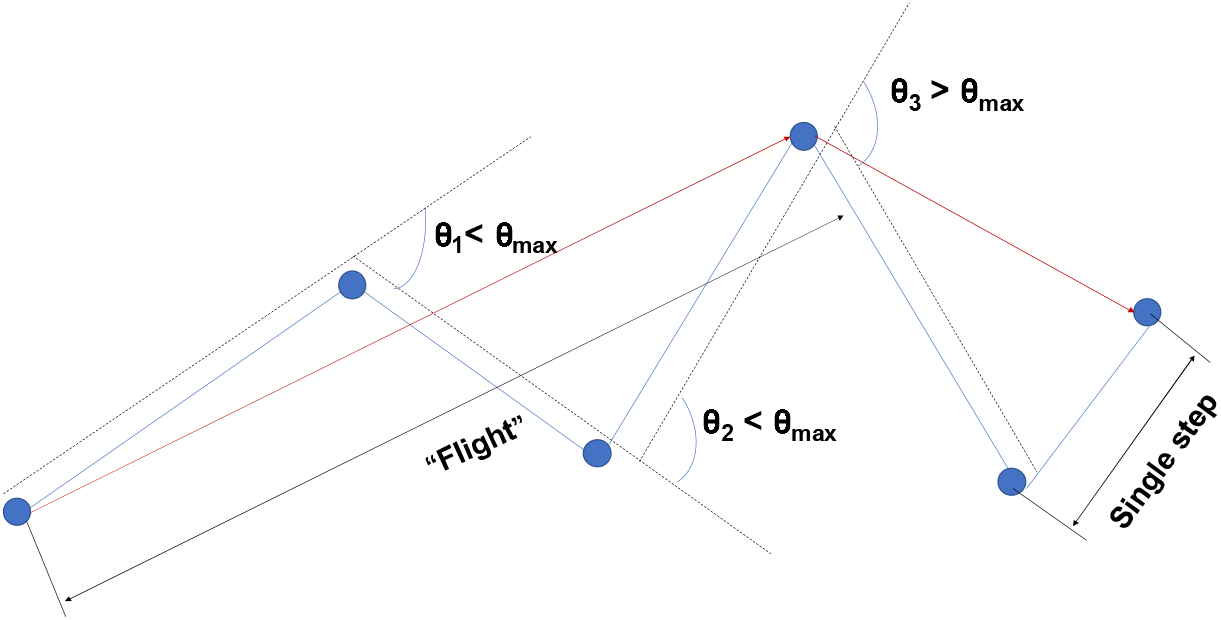

The long trajectory of the AuNR is segmented to several parts by the “angle method” modified from Turchin^1^, which is mainly based on the identification of the reorientation events during single particle walking. As illustrated in the figure above, multiple steps experienced in n time intervals are aggregated into a single “flight” segment, if the turning angle θ between the starting position and the end position is smaller than a predefined angle threshold value θ_max_. To determine the threshold value θ_max_, rather than stick to those reported in the literature, we test a wide range of angles from small to large and choose those angles that allow the resulting segment-lengths to converge to the power-law statistics (see discussions below). For the long single AuNR trajectories obtained in this study, it is discovered that whether the θ_max_ is too small (e.g. 5° or 10°) or too large (e.g. >100°), the “flight” segments are either over sampled or under sampled, leading to deviations during the fitting.

### Supplementary Note 2: Lévy walk models and the power law fitting

To characterize the power law statistics of the Lévy walk model, we used a continuous distribution^2,3^,

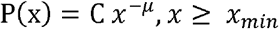

where α is a constant parameter of the distribution
 known as the exponent or scaling parameter. The scaling parameter typically lies in the range 1 < *μ* ≤ 3, although there are occasional exceptions. Since the distribution diverges at zero, so there must be a lower bound (*x_min_* > 0) on the power-law behavior. The approach uses log-likelihood fitting methods from the tail started at the lower bound *x_min_* that is determined in an automated way by a software developed by Aaron *et al*.^2^ For each possible choice of *x_min_*, the index value a is estimated via the method of maximum likelihood, and the Kolmogorov-Smirnov goodness-of-fit statistic D is calculated. The selected estimation of *x_min_* is the value that gives a minimum D over all values of *x_min_*. The selection of the power law model over the exponential model is justified using the Akaike weights that calculated the relative likelihoods of either model. See the detailed expression form and calculating methods in the reference article^4^.

### Supplementary Note 3: The auto-correlation function of velocity for LWs

Let v(t) denotes the velocity at time *t* of a particle following a LW. In the original LW model^5^, particles move at constant speed between random reorientations. We defined the velocity auto-correlation function^6^ as

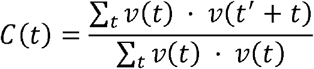

where 〈·〉 denotes averaging over all times *t* in independent samples of infinite trajectories. Assuming waiting times between reorientation events have a density ψ(*τ*) with a power-law tail, ψ(*τ*) ~ *τ*^−*β*+1^, then we have C(Δ*t*)~*t*^−(*β*−1)3,5,7^. Here, α=1.38, β = 3 − α =1.62 (*β* is the Levy stable parameter with α + β = 3). Thus, the C(t) should have a tail with about 0.6. We note that because of the “jump-and-linger” two-state motion of the AuNRs, it is normal to have a small deviation.

## Supplementary movies

**Supplementary Movie 1.** Individual AuNRs move on the upper layer of the swarming bacteria. The single AuNRs appear as red points. The scale bar is 15 μm.

**Supplementary Movie 2.** Trajectories of the single particle tracking results of multiple AuNRs. The scale bar is 10 μm.

**Supplementary Movie 3.** Polystyrene microbeads (0.5 μm) adhere on the air/liquid surface of the fluid layer of swarming bacteria.

**Supplementary Movie 4.** Gold nanospheres (120 nm) sink into the bottom of the bacteria fluid layer, rocking and colliding with the bacteria locally in a small area.

**Supplementary Movie 5.** AuNRs move in a Brownian way in the fluid layer after the bacteria have been exposed to the UV light illumination for a long period of time and stopped swarming.

**Supplementary Movie 6.** Particle image velocity (PIV) vectors overlaid on the raw image. The velocity vectors are shown in green arrows and the interpolations are shown in orange. The yellow circle marks the position of the AuNR being carried across the field. The moving single bacterium appears in long brown rod. The scale bar is 10 μm.

**Supplementary Movie 7.** PIV mapping overlaid on the PIV vectors, where the clockwise spin is shown in blue shades and the counter clockwise spin are shown in red shades. The scale bar is 10 μm.

## Supplementary Figures

**Supplementary Figure 1.**
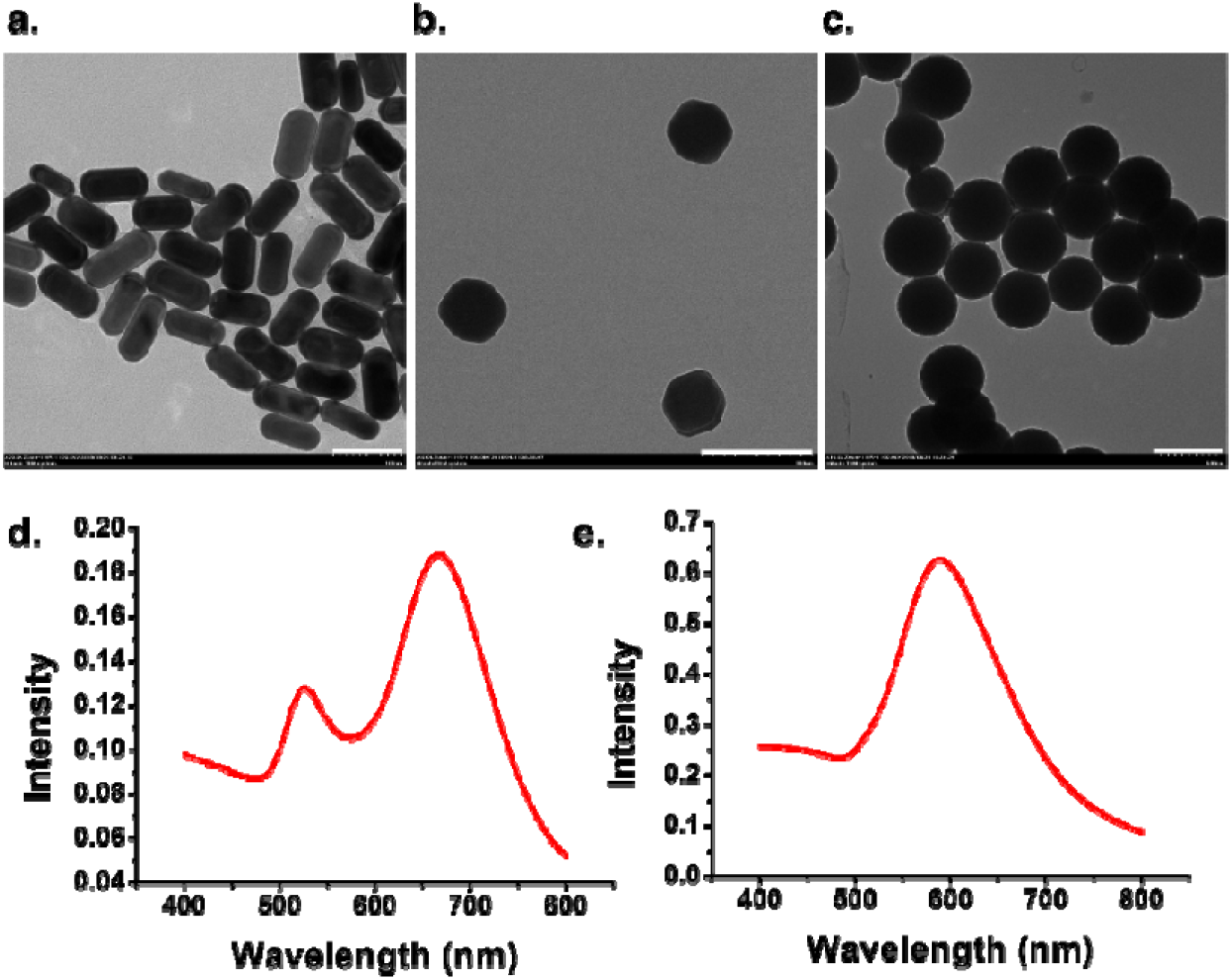
Characterization of the particles used in the experiment. (a) TEM images of 40×84 nm gold nanorods (AuNRs). The scale bar is 100 nm. (b) TEM images of 120 nm gold nanospheres. The scale bar is 200 nm. (c) TEM images of 0.5 μm polystyrene spheres. The scale bar is 500 nm. (d) UV-vis extinction spectra of the 40×84 nm AuNRs in (a). The longitudinal is 663 nm. (e) UV-vis extinction spectra of the 120 nm gold nanospheres in (b). The peak is 605 nm.

**Supplementary Figure 2.**
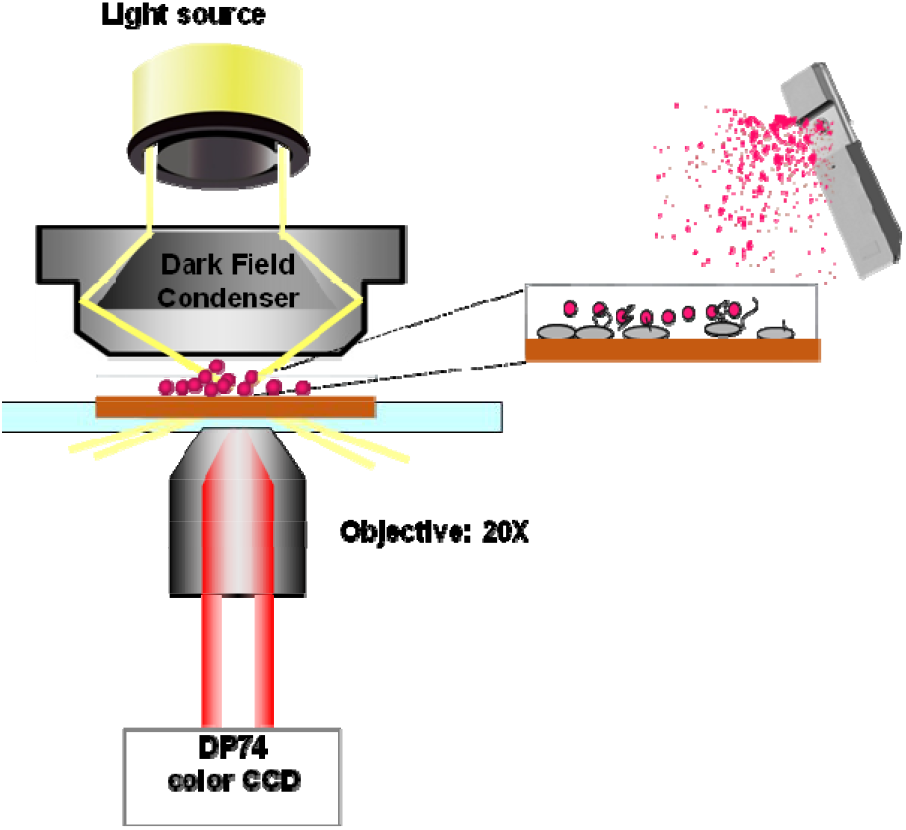
Schematic diagram of the imaging device. An inverted Nikon Ti-U microscope equipped with a 100 W halogen tungsten lamp, a dark-field air condenser, a 20X long working distance objective lens and an Olympus DP74 color CMOS camera is used to observe the swarming bacteria colony cultured in a 120 mm petri dish. The AuNRs are sprayed into the experimental system.

**Supplementary Figure 3.**
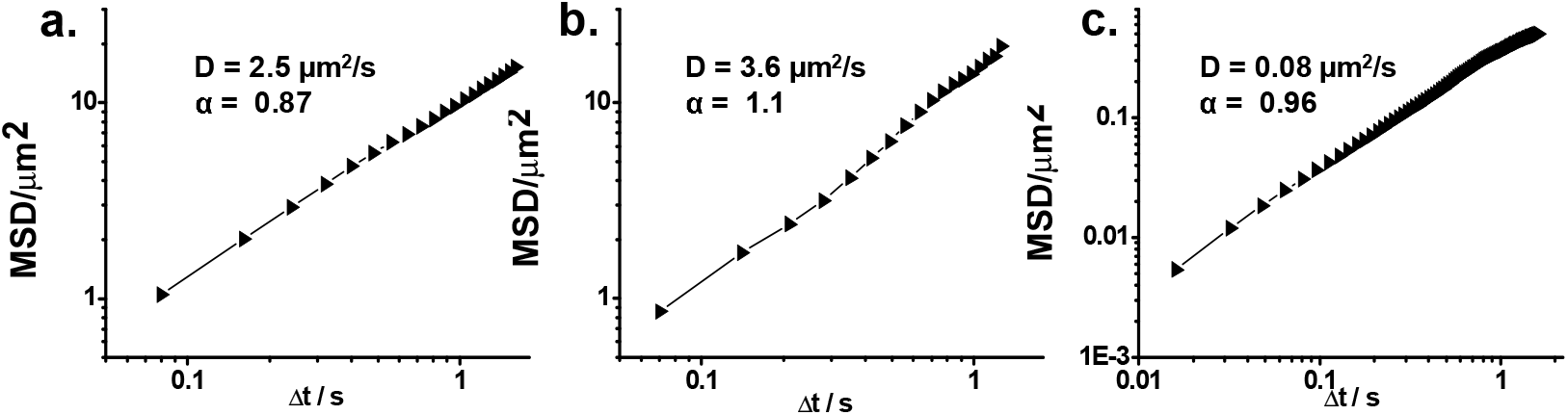
Ensemble-averaged mean square displacements (MSDs) of the control experiments. (a) MSDs of the AuNR movements in the LB cultured fluid medium where the motile bacteria are filtered out with a slope of 0.87 and a diffusion coefficient of 2.5 μm^2^/s. (b) MSDs of the AuNR movements in the LB culturing suspensions where the cells are swimming with a slope of 1.1 and a diffusion coefficient of 3.6 μm^2^/s. (c) MSDs of the AuNR movements in the fluid above the bacteria after the bacteria colony was irradiated under a UV lamp for 6 h with a slope α = 0.96 and a diffusion coefficient of 0.08 μm^2^/s. The MSD curves are calculated by averaging tens of trajectories.

